# Rapid Activation of Hematopoietic Stem Cells

**DOI:** 10.1101/2022.06.01.494258

**Authors:** Roshina Thapa, Erez Elfassy, Leonid Olender, Omri Sharabi, Roi Gazit

## Abstract

Adult Hematopoietic Stem Cells (HSCs) in the bone marrow (BM) are quiescent. Following perturbations, such as blood loss or infection, HSCs may undergo activation to accelerate the production of needed effector blood and immune cells. Surprisingly, little is known about the earliest stages of HSC activation. We utilize surface markers of HSC activation, CD69, and CD317, revealing response as early as 2 h after stimulation. The dynamic expression of HSC activation markers varies between viral-like (poly-Inosinic-poly-Cytidylic) or bacterial-like (Lipopolysaccharide) immune stimuli. We further quantify the dose response, demonstrating a low-threshold and similar sensitivity of HSCs and progenitors in the BM. Finally, we find a positive correlation between the expression of surface activation markers and early exit from quiescence into proliferation. Our data show that the premier response of adult stem cells to immune stimulation is rapid, sensitive, and directly leads to proliferation.

## Introduction

Hematopoietic Stem Cells (HSCs) are the source of all blood and immune cells (Sawai et al., 2016; Yamamoto et al., 2018). HSCs are largely quiescent in the adult bone marrow (BM), preserving long-term multipotency (Busch et al., 2015; Dzierzak and Speck, 2008; van der Wath et al., 2009; Yamazaki et al., 2007). When the need emerges, HSCs may undergo activation, and contribute to the accelerated hematopoiesis (Khosravi et al., 2014; Takizawa et al., 2011). The speed, as well as the sensitivity threshold of HSC activation, are yet to be determined.

Poly-Inosinic-poly-Cytidylic (pIpC) and Lipopolysaccharide (LPS) are the most commonly used viral- and bacterial-like immune stimuli, respectively (Chavakis et al., 2019). pIpC stimulation activates type-I interferon response, leading to the activation of HSCs in the BM (Essers et al., 2009; Sato et al., 2009). Such HSC activation depends largely on IFNα (Essers et al., 2011; Essers et al., 2009). Interestingly, pIpC was also reported to skew HSC differentiation into megakaryocytes (Haas et al., 2015), presumably to enhance coagulation. Moreover, chronic pIpC stimulation was shown to drive DNA damage and possible exhaustion of genetically susceptible HSCs (Walter et al., 2015), while chronic viral infection impairs HSC’s potency indirectly via CD8 T cells that change the BM niche (Isringhausen et al., 2021). LPS stimulate the major TLR4 pathway and activates HSCs directly and indirectly (Takizawa et al., 2017; Takizawa et al., 2011). Overwhelming stimulation can exhaust HSCs (Baldridge et al., 2010; de Bruin et al., 2013; Matatall et al., 2016; Pietras et al., 2016); however, milder models suggest that HSCs may re-enter quiescence and preserve potency despite prolonged stimulation (Pietras et al., 2014). Interestingly, LPS and related cytokines such as IL-1 and TNFα can skew differentiation into the myeloid rather than the lymphoid branch (Pietras et al., 2016; Takizawa et al., 2017; Yamashita and Passegue, 2019). Immune stimulation was further reported to induce trained immunity in HSCs, including metabolic and epigenetic myeloid skewing (de Laval et al., 2020; Mitroulis et al., 2018). Either viral- or bacterial stimuli induce the quiescent HSCs to proliferate – and give rise to more effector cells (Essers et al., 2011; Pietras et al., 2014; Takizawa et al., 2017; Takizawa et al., 2011; Walter et al., 2015; Yanez et al., 2009). Surprisingly, previous studies focused primarily on phenotypes emerging within days and months, but early stages of HSC activation haven’t been studied extensively.

We identified HSC-activation markers (Bujanover et al., 2018), including CD69 (*Clec2c*) and CD317 (*Bst2*). These markers were validated in a brilliant study showing the essential role of CD317 for retaining activated HSCs in the proper BM niche (Florez et al., 2020). In the current work, we sought to utilize these surface markers to study how early HSC activation occurs and how sensitive HSCs are to commonly used immune stimuli. Our quantified data show rapid response and high sensitivity, with a positive correlation between surface markers and the proliferative state of HSCs in the BM.

## Results

### Immune activation of hematopoietic stem and progenitor cells is rapid

We previously showed that HSC activation markers CD69 and CD317 are expressed at 24 h following immune stimulation (Bujanover et al., 2018). However, we did not find data on HSC response at earlier time points. Refining the time scale analysis of HSCs activation may reveal how fast they respond to the immune stimuli. Therefore, we stimulated mice with pIpC (100 µg, intra-peritoneal) and collected BM for FACS analysis using conventional HSC markers (see methods) plus CD69 and CD317 at 2, 4, and 24 h (Figure 1A). We observed an elevation of CD69 expression to more than 30% of the Lineage^-^Sca1^+^cKit^+^ compartment (LSK, encompassing multipotent stem- and progenitors), and in the LSKCD48^-^CD150^+^ HSC population after 2 h (Figure 1B). CD69 expression further increased at 4 and 24 h (Figures 1C and1D). Intriguingly, CD317 expression seemed to increase slightly slower, as seen at 2 and 4 h, with a peak at 24 h (Figures 1C and 1D). The BM progenitor populations showed a similar trend, changes in the CD317 expression on Lin^-^ or LK populations occurred with a comparable time-dependent scale, but the magnitude of activation was noticeably different (Figure S1). The more primitive populations, such as LSK and LSKCD48^-^150^+^ HSC, are far more sensitive to immune stimulation. In the case of pIpC stimulation, after 24 h, more than 80% of LSKCD48^-^CD150^+^ HSC and almost 100% of LSK cells elevated their CD317 expression. On the other hand, less the 60% of Lin^-^ and less than 80% of Lineage^-^cKit^+^ (LK) cells expressed CD317 at the same time point (Figures S1A-S1C). Of note, pIpC stimulation caused an elevation of Sca-1 expression, shifting more progenitors into the LSK gate 24 h after stimulation. pIpC mimics viral infection, we also wanted to examine a model for bacterial infection. LPS activates TLR4 and can activate HSCs both directly and indirectly. We used a dose of 20 µg, injected intra-peritoneal (Figure 1E). With this stimulation, CD69 expression peaked at 2 h, both in LSK and in the LSKCD48^-^CD150^+^ HSC populations, gaining about 90% positive cells (Figure 1G). However, with LPS stimulation, CD317 reached similar expression levels only at 4 h. Later, CD69 expression decreased, whereas CD317 levels were maintained in most cells (Figures 1F-1H). Furthermore, the elevation of CD69 and CD317 expression in other populations in the BM of the same mice was substantial as well (Figures S1D-S1F). Lin^-^ population, again, followed the same time-dependent trend in expression of both CD69 and CD317, peaking at 2 and 4 h after stimulation, respectively, but to a level of about 40% of cells (Figures S1D and S1E), followed by a decrease in expression in agreement with previously published data (Bujanover et al., 2018). In the case of the LK population, the baseline of CD69 expression was 25% of cells, with a peak of about 60% 4 h after the immune stimulation. CD317 expression elevated after 4 h to about 90% of the cells. The expression of both CD69 and CD317 decreased after 24 h, with CD69 levels returning to baseline levels and CD317 remaining at about 60% (Figures S1D and S1F). Notably, LPS stimulation, like pIpC stimulation, caused an elevation of Sca-1 expression, suggesting that some of the cells with the LSK immune phenotype might have gained it as a direct result of the immune stimulation. These findings suggest CD69 to be more responsive during the early phase of immune stimulation (2 h), whereas CD317 elevated at the later phase of immune stimulation (24 h). Our data also suggests higher sensitivity of more primitive hematopoietic cell populations to the introduction of viral or bacterial immune stimuli.

**Figure 1.**
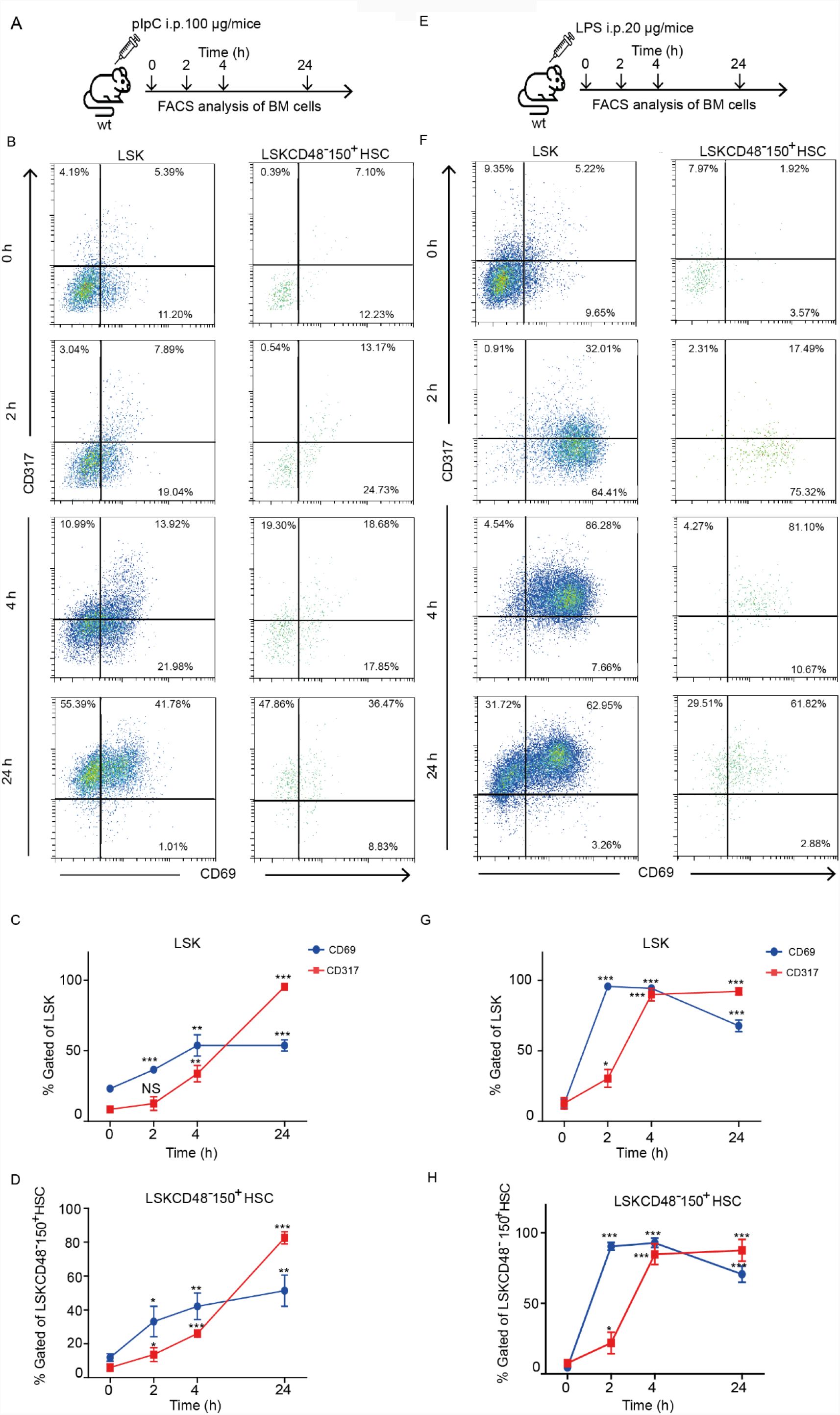
Immune activation of hematopoietic stem and progenitor cells is rapid. (A) Schematic representation of the experimental settings involving injection of pIpC at different time points (0, 2, 4, and 24 h). (B) Representative FACS plots showing the percentage of CD69^+^ and CD317^+^ cells in the LSK (left panels) and LSKCD48^-^150^+^ HSC (right panels) populations from control and pIpC stimulated mice at different time points. (C) and (D) Quantification of CD69 and CD317 expression in the LSK (C) and LSKCD48^-^150^+^ HSC populations (D) in the BM of control and pIpC stimulated mice. (E) Schematic representation of the experimental settings involving injection of LPS at different time points (0, 2, 4, and 24 h). (F) Representative FACS plots showing the percentage of CD69^+^ and CD317^+^ cells in the LSK (left panels) and LSKCD48^-^150^+^ HSC (right panels) populations from control and LPS-stimulated mice at different time points. (G) and (H) Quantification of CD69 and CD317 expression in the LSK(G) and LSKCD48^-^150^+^ HSC (H) populations in the BM of control and LPS-stimulated mice. Data are presented as average ± SD, a summary of three independent experiments, n≥3 mice per group; * - p<0.05, ** - p<0.01, *** - p<0.001.

### CD317 reveals a dose-response to pIpC above a minimal threshold

Following the findings of rapid response to pIpC and accumulation of CD317 at 24 h, we next wanted to examine if HSCs activation is dose-dependent or rather an all-or-none response. To test this hypothesis, we challenged mice with various doses of pIpC, ranging from 0.01 µg to 100 µg, and analyzed the expression of CD317 in different populations of BM hematopoietic cells. The CD317 expression in the broader LSK and the LSKCD48^-^CD150^+^ HSC was proportional to an increasing dose of pIpC, indicating 0.1 µg of pIpC as the minimal dose for the induction of CD317 expression. An increase in the dose of pIpC led to an increase in the expression of CD317 to 80% (Figures 2A-2C). The minimal pIpC dose from our data was further endorsed by the EC50 calculation: 0.10 µg for LSK and 0.12 µg for LSKCD48^-^CD150^+^ HSC Furthermore, the relationship between pIpC dose and CD317 expression fitted well with linear regression (Figures 2D and 2E). Similarly, Lin^-^ or LK cells followed a dose-response trend in CD317 expression with pIpC (Figure S2). However, the maximal increment observed within the Lin^-^ population was much lower (expression increased to a maximum of 40% with the highest dose of pIpC) than that observed in LSK cells and the LSKCD48^-^CD150^+^ HSC (expression increased to a maximum of 80% with the highest dose of pIpC, Figures 2 and S2). Notably, the LK cell population showed a stronger response than the Lin^-^ population, with CD317 expression reaching around 60% (Figure S2). Furthermore, calculations of the EC50 values for the CD317 activation in Lin^-^ and LK populations also fitted well with linear regression and were found to be 0.07 µg and 1.30 µg, respectively (Figure S2D and S2E). Therefore, hematopoietic stem- and progenitor cells have a dose-dependent response to pIpC in terms of surface expression of CD317 throughout a broad range of concentrations.

**Figure 2.**
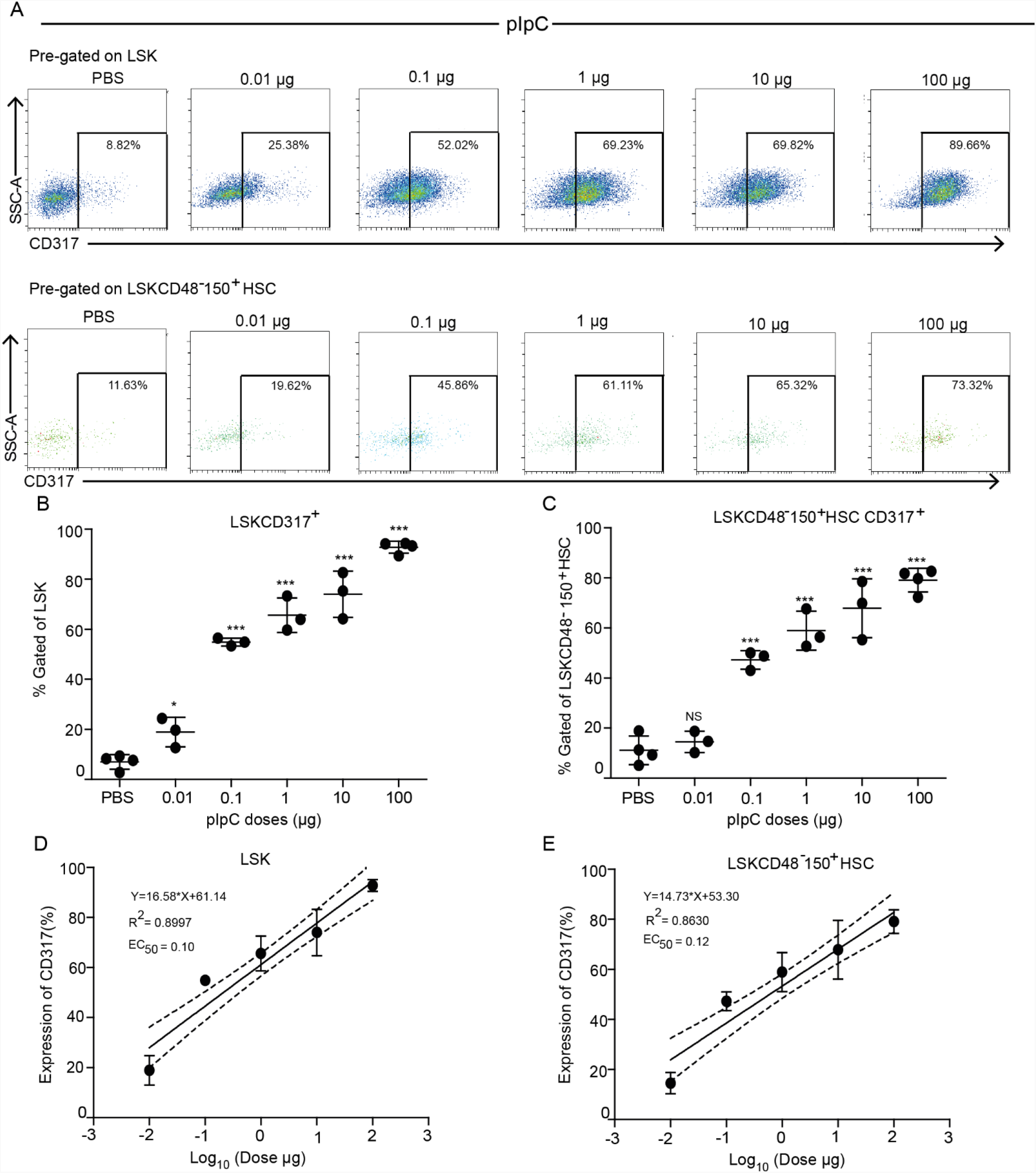
CD317 reveals a dose-response to pIpC above a minimal threshold. (A) Representative FACS plots showing the percentage of CD317^+^ cells in the LSK (upper panels) and LSKCD48^-^150^+^ HSC (lower panels) populations from the BM of PBS-treated (control) and pIpC-stimulated mice under various pIpC doses (0.01 µg -100 µg as positive control) at 24 h post-injection. (B) and (C) Quantification of the LSK CD317^+^ (B) and LSKCD48^-^150^+^ HSC CD317^+^ (C) cell populations from the BM of PBS-treated (control) mice and mice treated with PBS (control) or different doses of pIpC. (D) and (E) Linear regression model fitting the dose-dependent response with the expression of CD317 in LSK (D) and LSKCD48^-^150^+^ HSC (E) populations. Solid lines represent the linear fit of data. Dotted lines represent 95% confidence intervals. Data are presented as average ± SD, a summary of three independent experiments, n≥3 mice per group; * - p<0.05, ** - p<0.01, *** - p<0.001.

### CD69 elevation is dose-dependent at higher concentrations of LPS stimulation

Next, following the robust response described above (Figure 1), we examined the dose-dependent effect of LPS on the expression of CD69. We challenged mice with various doses of LPS (0.0001-20 µg) and analyzed the expression of CD69 in different hematopoietic cell populations in the BM 2 h later. The CD69 expression in the LSK and the LSKCD48^-^CD150^+^ HSC populations was significantly upregulated at relatively higher doses of LPS (0.1 µg onwards), with maximal upregulation observed at 1 µg of LPS More specifically, in the LSK population, 50% of cells expressed CD69 at 0.1 µg of LPS, with expression levels reaching 80% with 1 µg of LPS (Figures 3A and 3B). Similarly, CD69 expression in LSKCD48^-^CD150^+^ HSC population increased from 25% to 60% at similar LPS doses (0.1 µg and 1 µg, respectively, (Figures 3A and 3C). The EC50 value of 0.07 µg (LSK), and 0.21 µg (LSKCD48^-^ CD150^+^ HSC), delineated a threshold for LPS on CD69 elevation in these populations. Nevertheless, we gained a clear dose response in terms of CD69 expression with the higher LPS concentrations using the linear regression tool (Figure 3D and3E). The threshold for LPS was even more pronounced in the Lin^-^ and LK populations. such that only 1 µg of LPS was able to exert a significant effect on the expression of CD69, with the overall increment reaching up to 20% with 20 µg of LPS (Figures S3A-S3C). Despite having a lower maximal activation in the Lin^-^ or LK, the EC50 values of 0.39 µg (Lin^-^), and 0.17 µg (LK) are in the same range as LSK or HSCs (Figures S3D and S3E).

**Figure 3.**
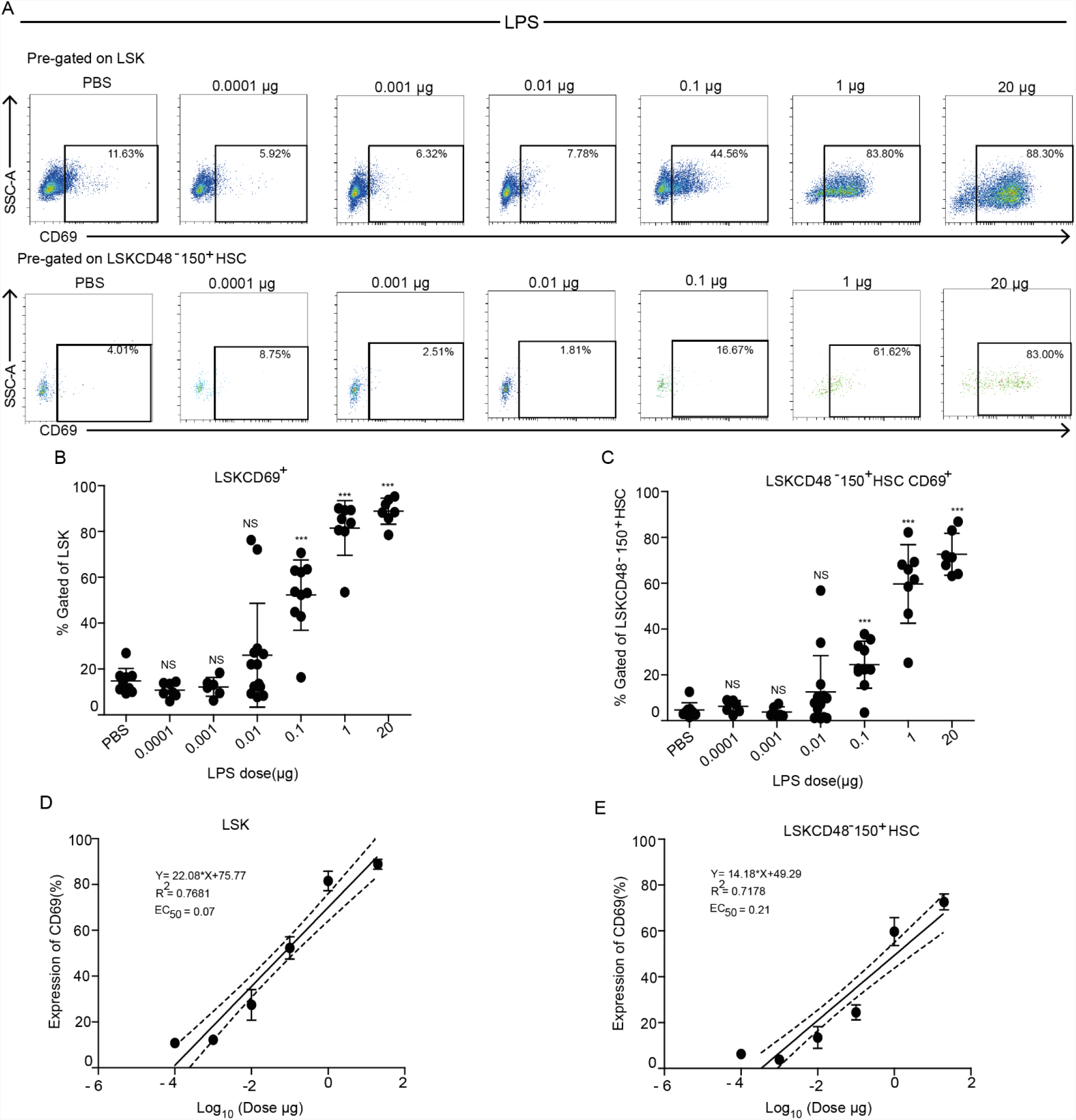
CD69 elevation is dose-dependent at higher concentrations of LPS. (A) Representative FACS plots showing the percentage of CD69^+^ cells in the LSK (upper panels) and LSKCD48^-^150^+^ HSC (lower panels) populations from the BM of PBS-treated (control) mice and mice stimulated with various doses of LPS (0.0001-20 µg) at 2 h post-injection. (B) and (C) Quantification of the LSK CD69^+^ (B) and LSKCD48^-^150^+^ HSC CD69^+^ (C) cell populations from the BM of PBS-treated (control) mice and mice treated with PBS (control) or different doses of LPS. (D) and (E) Linear regression model fitting the dose-dependent response with the expression of CD69 in LSK (D) and LSKCD48^-^150^+^ HSC (E) populations. Solid lines represent the linear fit of data. Dotted lines represent 95% confidence intervals. Data are presented as average ± SD, a summary of three independent experiments, n≥3 mice per group; * - p<0.05, ** - p<0.01, *** - p<0.001.

### CD69 and CD317 expression in activated stem and progenitor populations positively correlates with proliferation

To evaluate whether elevation in activation markers correlates with hematopoietic stem and progenitor cell proliferation, we conducted a cell cycle analysis. In this set of experiments, we tested for Ki67 expression in pIpC- and LPS-stimulated mice at 4 and 24 h or 2 and 24 h, respectively (Figures 4A and 4D). At designated time points, we extracted the BM, stained for surface markers, fixed and permeabilized the cells, then stained for the nuclear Ki67 and DNA-content (see methods). Surprisingly, at 4 h post pIpC stimulation, a significant proportion of LSK CD317^+^ cells, but not LSK CD317^-^ cells, were in the S and M phase of the cell cycle (Figure 4B). The CD317^+^ fractions of LSK and HSC populations had fewer cells in the G0 phase, and this trend was sustained at 24 h (Figures 4B and 4C). Similar differences between division kinetics of CD317^+^ and CD317^-^ cells were also observed for Lin^-^ and LK populations (Figures S4B and S4C). We also tested the correlation between CD69 expression and proliferation following stimulation with LPS. Here only a slight change was observed in the LSK population at 2 h, reaching significance at 24 h, with CD69^+^ being less quiescent and more proliferating (Figure 4E). Surprisingly, we observed more G0 cells in the CD69^+^ fraction than in their CD69^-^ counterparts within the LSKCD48^-^CD150^+^ HSC compartment 2 h after the stimulation (Figure 4F). At 24 h, the CD69^+^ HSCs were not more proliferating than the CD69^-^ HSCs, and their respective G0 fractions were almost equal (Figure 4F). As for the Lin^-^ and LK populations, their CD69^+^ fractions exhibited more proliferation than CD69^-^ 2 h after the stimulation, but these differences were largely diminished at 24 h (Figure S4E-S4F). Taken together, with one peculiar exception of HSCs at the early 2 h time point, our data demonstrate the positive correlation of surface activation markers and proliferation of stem or progenitors following pIpC and LPS stimulation.

**Figure 4.**
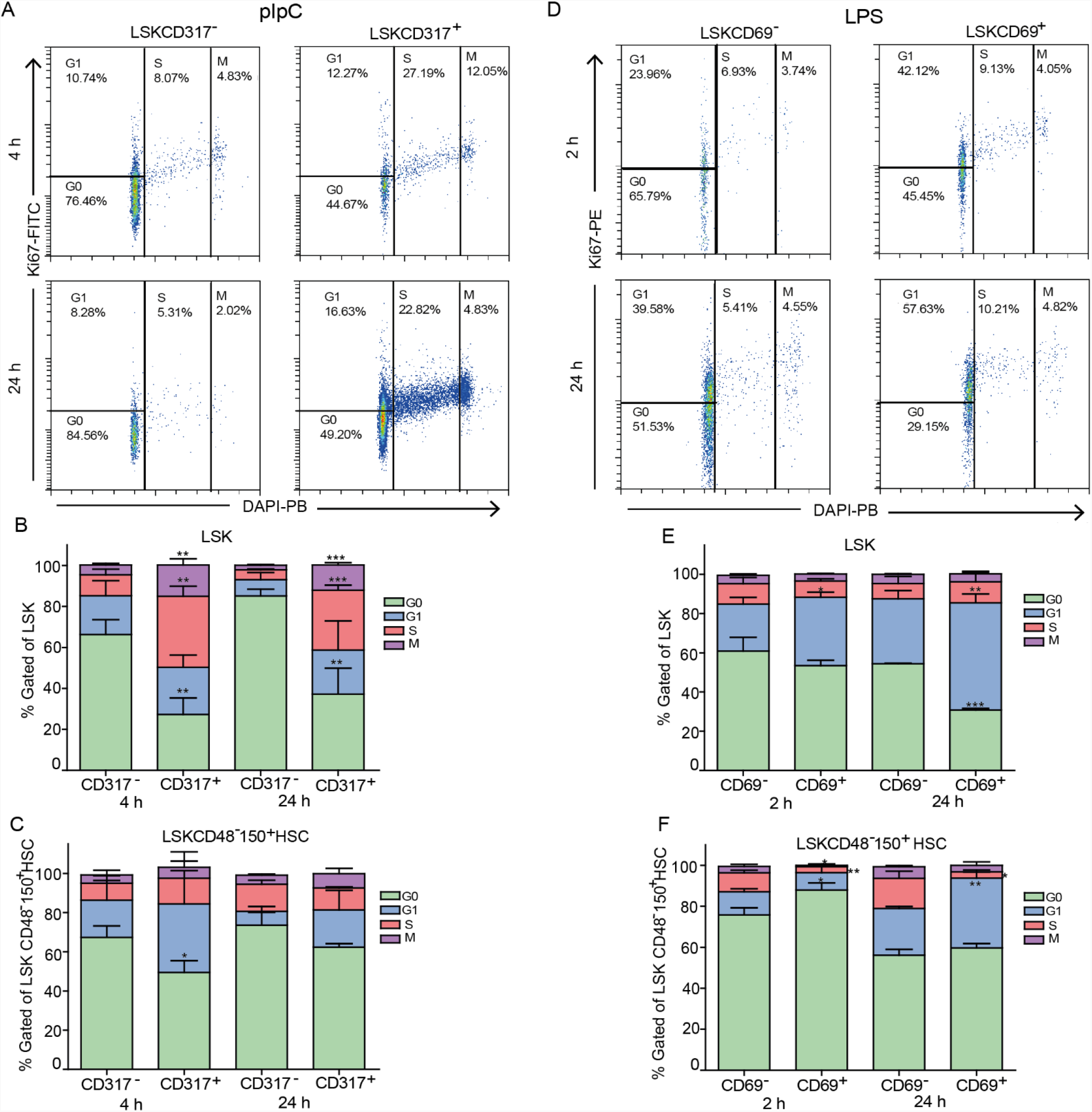
CD69 and CD317 expression on activated stem and progenitor populations positively correlate with proliferation. (A) Representative FACS plots showing cell cycle analysis using DAPI and intracellular expression of Ki67 in LSK CD317^-^ (left panel) and LSK CD317^+^ (right panel) cells from pIpC-stimulated mice at 4 and 24 h after injection. (B) and (C) Quantification of cell cycle phases (G0, G1, S, and M) in CD317^-^ and CD317^+^ fractions of the LSK (B) and of LSKCD48^-^150^+^ HSC populations from pIpC-stimulated mice at 4 h and 24 h post-injection. (D) Representative FACS plots showing cell cycle analysis using DAPI and intracellular expression of Ki67 in LSK CD69^-^ (left panel) and LSK CD69^+^ (right panel) cells from LPS-stimulated mice at 2 and 24 h after injection. (E) and (F) Quantification of cell cycle phases (G0, G1, S, and M) in CD69^-^ and CD69^+^ fractions of the LSK (E) and LSKCD48^-^ CD150^+^ HSC (F) populations from LPS-stimulated mice at 2 h and 24 h post-injection. Data are presented as average ± SD, a summary of three independent experiments, n≥3 mice per group; * - p<0.05, ** - p<0.01, *** - p<0.001.

## Discussion

This study presents the rapid activation of HSCs following immune activation in vivo. Utilizing surface markers, we demonstrate that HSCs gain rapid activation as early as 2 h after the stimulation. We demonstrate high sensitivity of primitive hematopoietic cells to pIpC and LPS, with a dose-dependent response range and a potentially low threshold. Finally, we indicate a correlation between the expression of the activation markers and proliferation in these cells, suggesting that the exit from quiescence is faster than previously estimated.

The immune response is rapid, showing pronounced elevation of systemic inflammatory cytokines by 2 h (Copeland et al., 2005). HSCs may sense immune activation both directly and indirectly, involving multiple receptors for microbial products (TLRs) and cytokines receptors for IL-1, IL-6, TNFα, and IFN (Chavakis et al., 2019; Clapes et al., 2016). As the crucial role of HSCs is to produce more hematopoietic cells, previous studies focused on their proliferation and differentiation following immune stimulation (Chavakis et al., 2019). Surface activation markers allow us to quantify earlier stages with a single-cell resolution (Figure 1). CD317 (BST2) has a role in anti-viral response (Le Tortorec et al., 2011), and it was also shown to play a role in the retention of HSCs within the BM (Florez et al., 2020). CD69 is a major activation marker of immune cells and a metabolic regulator (Cibrian and Sanchez-Madrid, 2017); intriguingly, CD69 is also implicated in HSC and Progenitors (HSPC) retention in the BM (Notario et al., 2018), suggesting that CD69, like CD317, may regulate their BM localization following stimulation.

Realizing the fast activation of HSCs positions them at the front line of the immune response, even at their BM-niche. A front line role for HSPCs was previously suggested in peripheral, secondary lymphoid tissues (Massberg et al., 2007). In this study, we focus on the BM, where most adult HSCs reside. Intriguingly, the sensitivity of HSCs is comparable to that of the hematopoietic progenitors in the BM (Figures 2 and 3) and other effector immune cells in vivo (Poast et al., 2002; Raveche and Steinberg, 1982; Tough et al., 1997), suggesting a coordinated systemic response. Thus, our data point HSCs to be an integral part of the immune system. HSCs are not a distal reservoir for late stage, but a proximal part of the acute immune response.

The activation of HSCs might have later consequences. Trained immunity, the adapted immunity of myeloid cells, suggests fine-tuning of response (de Laval et al., 2020). It will be most interesting to realize how deeply trained immunity is within the hematopoietic hierarchy. Our data suggest that HSCs do sense and gain rapid activation at a low threshold, providing possible “training” through multiple experiences throughout life. Intriguingly, the aging of HSCs is intrinsic (Rossi et al., 2005) but not yet characterized at the molecular level. HSCs were previously suggested to count cell divisions (Bernitz et al., 2016; Haas et al., 2018) – thus, proliferation must be tightly regulated. Indeed, we recently reported that HSCs might ignore an acute hypersensitivity immune response (Bujanover et al., 2021). Importantly, blocking inflammation was shown to attenuate the deleterious impact on HSPCs (Hernandez et al., 2019), providing strong support to prophylactic treatments. This study quantifies the rapid response of HSCs. It offers further options for prospective studies focused on modulations of activation in our long-term multipotent reservoir for blood and immune cells.

## Limitations

The prospective isolation of HSCs allowed major advancements, but some of the markers do change following immune stimulation, as we and others reported (Bujanover et al., 2018). A recent study suggests an improved protocol adding both CD34 and EPCR markers to better identify functional HSCs following immune stimulation (Rabe et al., 2020). We chose to use the common LSKCD48^-^CD150^+^ gating as we focus on early time points. For the same reason, we limited our study to BM and did not explore HSC mobilization, which may occur at later time points. Notably, highly-purified immune-phenotypic HSC population may consist of 1/3 functional long-term reconstituting cells, and these HSCs are heterogeneous (Haas et al., 2018). Our data show that virtually all the LSKCD48^-^CD150^+^ cells, encompassing all HSCs, gain activation markers. We also limit our experiments to key time points, reducing the number of animals. Future studies may fine-tune the time course and further examine possible heterogeneity among earlier- and later-activated HSCs and progenitors.

## Methods

### Key resource table

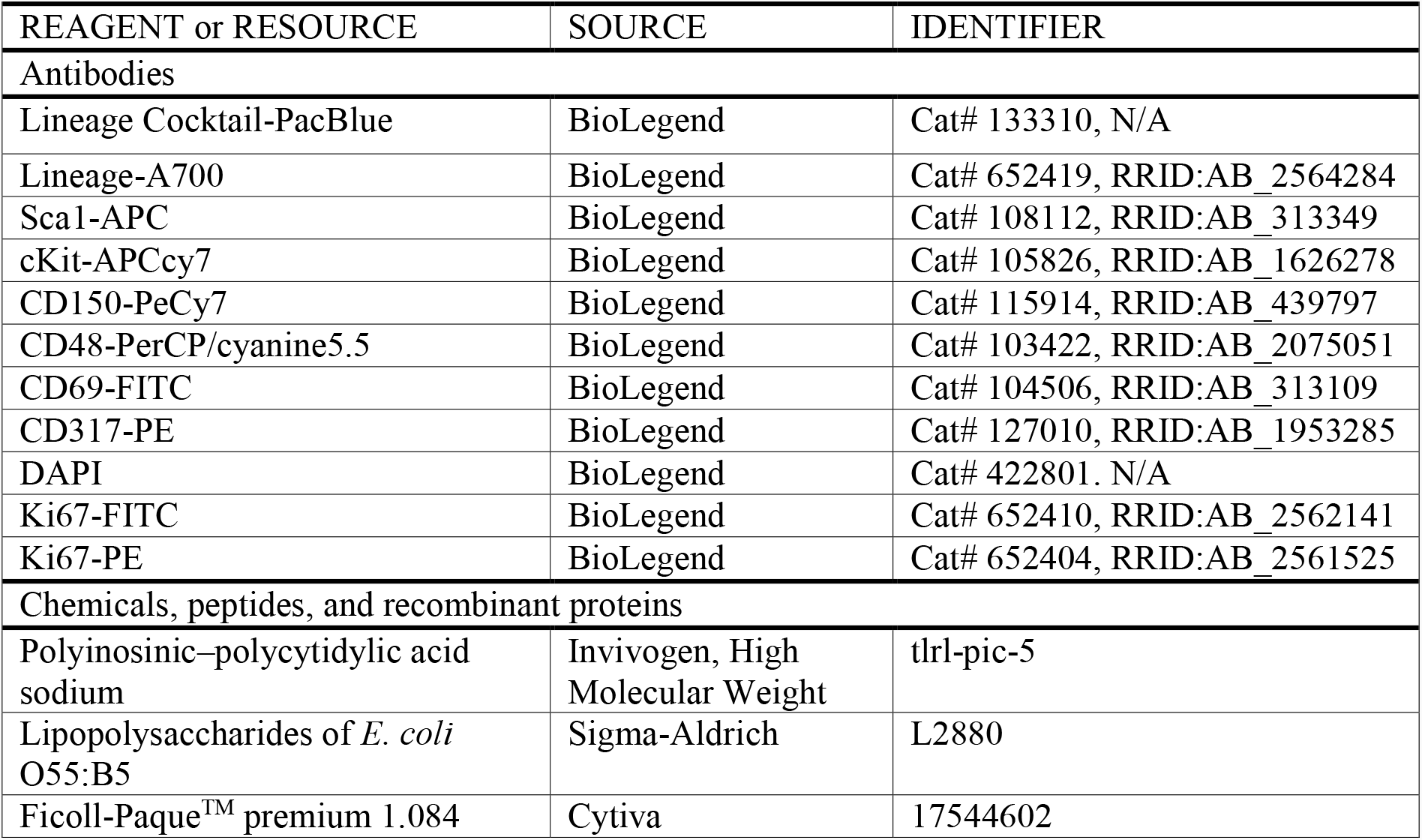
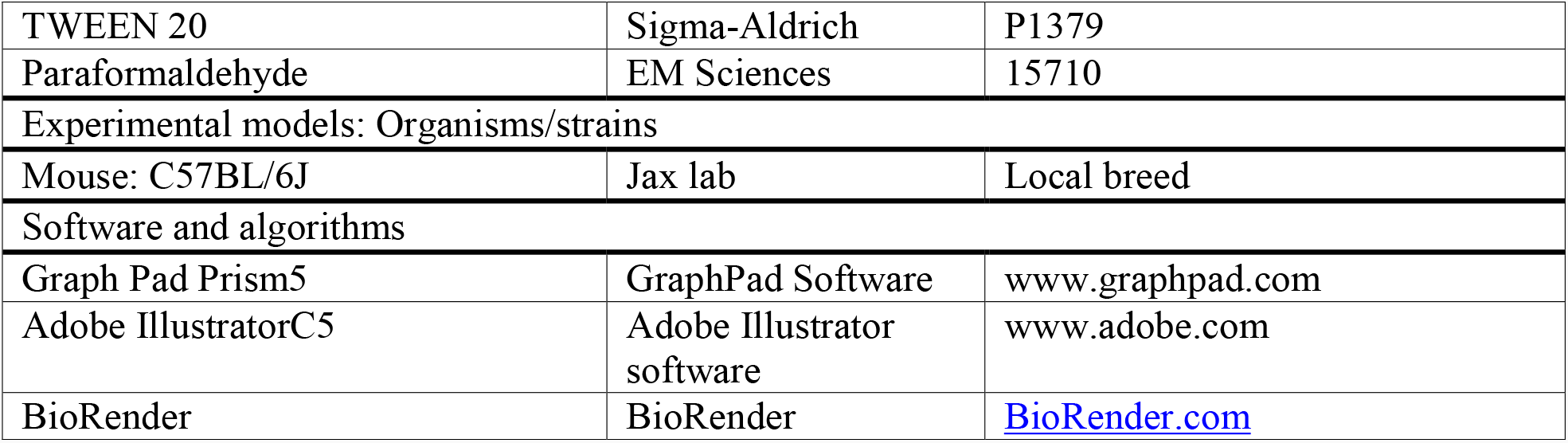

#### Mice and Ethical

All mice were kept at the Ben-Gurion University SPF (specific pathogen-free) unit. Mice strains used were C57Bl/6. Both male and female mice at 8-12 weeks old were used for the experiments, with age- and gender-matched littermates as control. All experiments were carried out according to the guidelines of the ethical committee and after approval of the local and state IACUC (IL-68-11-2018-D).

#### Immune stimulation

Mice were stimulated by poly-Inosinic–poly-Cytidylic acid sodium (pIpC: Invivogen, High Molecular Weight, catalog code: trl-pic-5) intraperitoneally (i.p.) 100 µg per mouse. Lipopolysaccharides of E. coli O55:B5 (LPS, Sigma-Aldrich cat: L2880 lot: 113M4068V) were REAGENT or RESOURCE SOURCE IDENTIFIER Antibodies Lineage Cocktail-PacBlue BioLegend Cat# 133310, N/A Lineage-A700 BioLegend Cat# 652419, RRID:AB_2564284 Sca1-APC BioLegend Cat# 108112, RRID:AB_313349 cKit-APCcy7 BioLegend Cat# 105826, RRID:AB_1626278 CD150-PeCy7 BioLegend Cat# 115914, RRID:AB_439797 CD48-PerCP/cyanine5.5 BioLegend Cat# 103422, RRID:AB_2075051 CD69-FITC BioLegend Cat# 104506, RRID:AB_313109 CD317-PE BioLegend Cat# 127010, RRID:AB_1953285 DAPI BioLegend Cat# 422801. N/A Ki67-FITC BioLegend Cat# 652410, RRID:AB_2562141 Ki67-PE BioLegend Cat# 652404, RRID:AB_2561525 Chemicals, peptides, and recombinant proteins Polyinosinic–polycytidylic acid sodium Invivogen, High Molecular Weight tlrl-pic-5 Lipopolysaccharides of E. coli O55:B5 Sigma-Aldrich L2880 Ficoll-PaqueTM premium 1.084 Cytiva 17544602 TWEEN 20 Sigma-Aldrich P1379 Paraformaldehyde EM Sciences 15710 Experimental models: Organisms/strains Mouse: C57BL/6J Jax lab Local breed Software and algorithms Graph Pad Prism5 GraphPad Software www.graphpad.com Adobe IllustratorC5 Adobe Illustrator software www.adobe.com BioRender BioRender BioRender.com administered i.p. at 20 µg per mouse. Stimulation was done at various time points (2, 4, and 24 h). Unstimulated mice were used as control, such as time 0. For dose-response, mice were stimulated by Polyinosinic–polycytidylic acid sodium salt (pIpC: Invivogen, High Molecular Weight, catalog code: tlrl-pic-5) intraperitoneally (i.p.) with various doses (0.01, 0.1, 1, and 10 µg), and analyzed at 24 h. Unstimulated mice were used as control. Lipopolysaccharides of E. coli O55:B5 (LPS, Sigma-Aldrich cat: L2880 lot: 113M4068V) were administered i.p. with various doses (0.0001, 0.001, 0.01, 0.1, and 1 µg) per mouse, and analyzed at 2 h.

#### Flow cytometry

BM cells were extracted from the tibia, femur, and pelvis using mortar and pestle; sample media comprised of Phosphate Buffer Saline (PBS) with 2mM EDTA and 2% Foetal Bovine Serum (FBS Lot#2137767, Biological Industries). Mononuclear cells were enriched over Ficoll-PaqueTM premium 1.084 (Cytiva, cat #17544602) and stained using the following antibodies (all from Biolegend): Lineage Cocktail-PacBlue, Lineage-A700, Sca1-APC, cKit-APCcy7, CD150-PeCy7, CD48-PerCP/cyanine5.5, CD69-FITC, CD317-PE, and DAPI. FACS Cytoflex LX (Beckman Coulter) and FacsAriaIII (BD) were used for analysis and sorting. Cytexpert software was used to analyze and visualize FACS data.

#### Cell cycle analysis

BM cells were extracted from the tibia, femur, and pelvis; mononuclear cells were enriched over histopaque and stained: Lineage-A700, Sca1-APC, cKit-APCCy7, CD150-PECy7, CD48-PC5.5, CD69-FITC, and CD317-PE. Cells were fixated in a 96U plate in 250 µl of 2% paraformaldehyde (PFA, EMS #15710) in PBS at room temperature for 20 min, washed twice in PBS, permeabilized in 0.25%Tween 20 (Sigma-Aldrich, P1379) in PBS for 30min, washed twice in 0.1% tween PBS then stained by Ki67-FITC or Ki67-PE (Biolegend) in 1:300 of stock in 0.1% tween PBS overnight. DAPI (10 µg/ml) was added to the cells before flow-cytometric analysis.

#### Statistics

FACS data are shown as mean ± SD. Data are representative of at least three independent experiments unless otherwise noted. A two-tailed T-test was performed, with p < 0.05 considered significant. Linear regression statistical tool was performed using the Graphpad prism.

## Acknowledgments

We thank all members of the R.G. laboratory for helpful discussions. Grant support MOSTDKFZ#CA179, and ISF#883/21.

## Contributions

EE and RT performed experiments and collected data; EE, RT, LO and OS analyzed and interpreted data; RG concept and designed the study; EE, RT, and RG wrote the manuscript.

## Supplemental Information for manuscript

**Figures S1-S4**

**Figure S1, Related to Figure 1.**
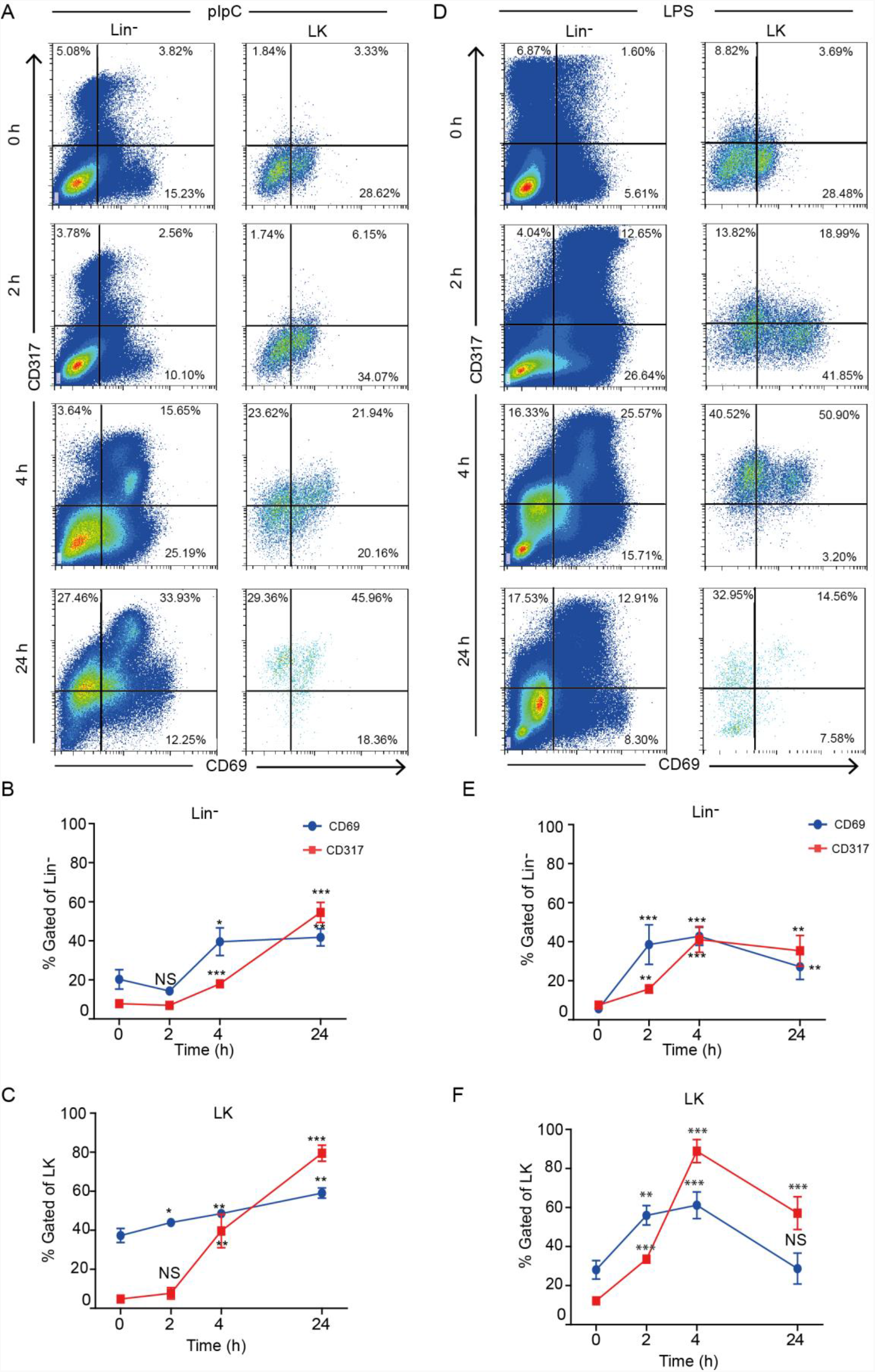
Early activation of hematopoietic progenitor cells upon immune stimulation. (A) Representative FACS plots showing the percentage of CD69^+^ and CD317^+^ in the Lineage^-^ (Lin^-^, left panels) and Lineage^-^ cKit^+^ (LK, right panels) populations from PBS-treated (control) and pIpC-stimulated mice at different time points. (B) and (C) Quantification of CD69 and CD317 surface expression in Lin^-^ (B) and LK (C) populations from the BM of pIpC-treated mice at different time points. (D) Representative FACS plots showing the percentage of CD69^+^ and CD317^+^ in the Lineage^-^ (Lin^-^, left panels) and Lineage^-^ cKit^+^ (LK, right panels) populations from PBS-treated (control) and LPS-stimulated mice at different time points. (E) and (F) Quantification of CD69 and CD317 surface expression in Lin^-^ (E) and LK (F) populations from the BM of LPS-treated mice at different time points. Data are presented as average ± SD, a summary of three independent experiments, n≥3 mice per group; * - p<0.05, ** - p<0.01, *** - p<0.001.

**Figure S2, Related to Figure 2.**
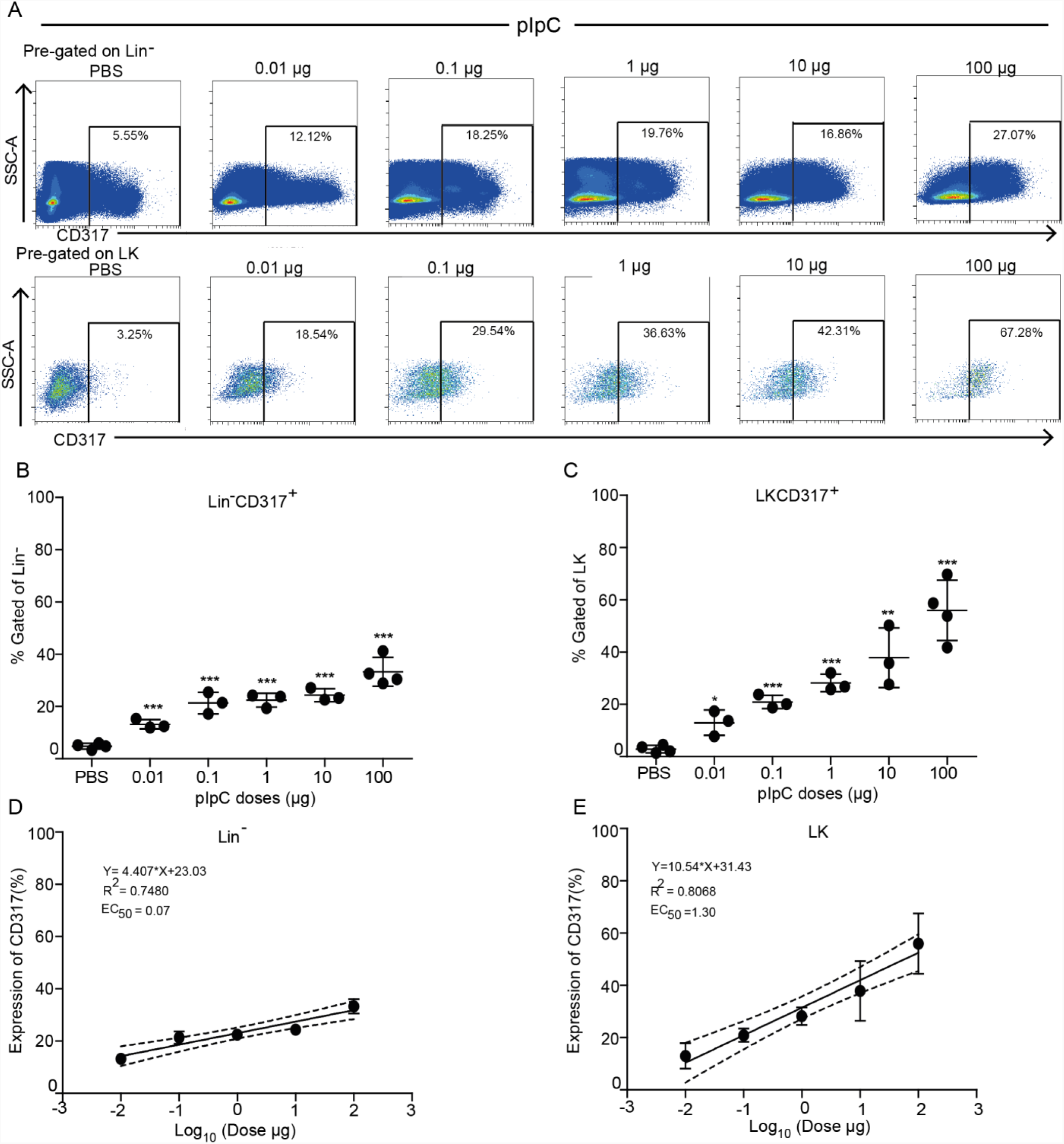
Dose-response of pIpC on activation of hematopoietic progenitor cells. (A) Representative FACS plots showing the percentage of CD317^+^ cells in the Lin^-^ (upper panels) and LK (lower panels) populations from the BM of PBS-treated (control) mice and mice treated with various doses of pIpC (0.01 µg -100 µg) at 24 h. (B) and (C) Quantification of CD317^+^ fractions in Lin^-^ (B) and LK (C) populations in the BM of mice treated with PBS (control) or various doses of pIpC. (D) and (E) Linear regression model fitting the dose-dependent response with the expression of CD317 in Lin^-^ (D) and LK (E) populations. Solid lines represent the linear fit of data. Dotted lines represent 95% confidence intervals. Data are presented as average ± SD, a summary of three independent experiments, n≥3 mice per group; * - p<0.05, ** - p<0.01, *** - p<0.001.

**Figure S3, Related to Figure 3.**
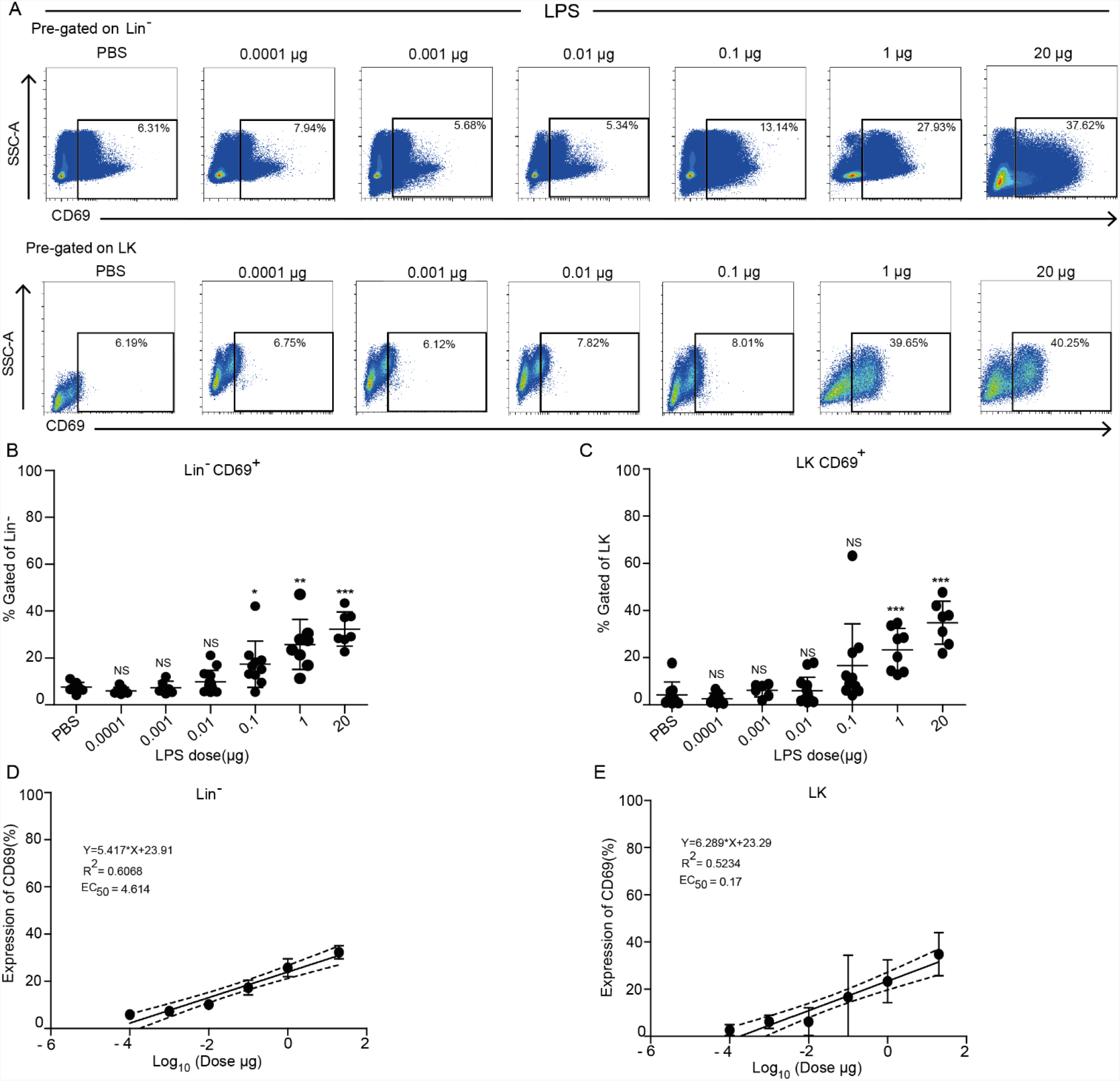
Dose-dependent activation of hematopoietic progenitor cells by LPS. (A) Representative FACS plots showing the percentage of CD69^+^ cells in the Lin^-^ (upper panels) and LK (lower panels) populations from the BM of PBS-treated (control) mice and mice treated with various doses of LPS (0.01 µg -100 µg) at 2 h. (B) and (C) Quantification of CD69^+^ fractions in Lin^-^ (B) and LK (C) populations in the BM of mice treated with PBS (control) or various doses of LPS. (D) and (E) Linear regression model fitting the dose-dependent response with the expression of CD69 in Lin^-^ (D) and LK (E) populations. Solid lines represent the linear fit of data. Dotted lines represent 95% confidence intervals. Data are presented as average ± SD, a summary of three independent experiments, n≥3 mice per group; * - p<0.05, ** - p<0.01, *** - p<0.001.

**Figure S4, Related to Figure 4.**
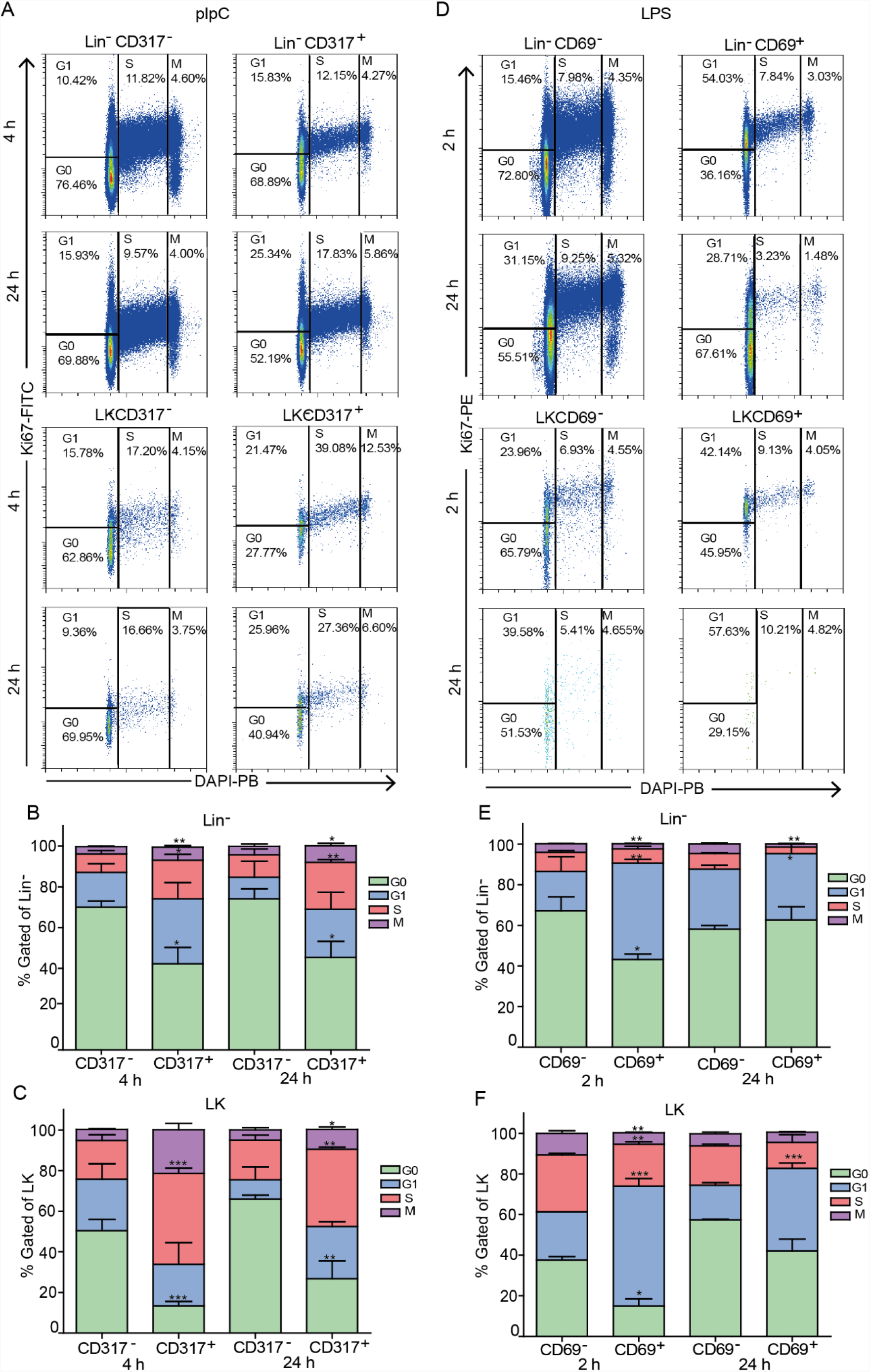
Activation of hematopoietic stem- and progenitor cells is linked with elevated proliferation. (A) Representative FACS plots showing cell cycle analysis using DAPI and intracellular expression of Ki67 in CD317^-^ and CD317^+^ fractions of Lin^-^ (upper panel) and LK (lower panel) cell populations from the BM of pIpC-stimulated mice at 4 h and 24 h post-injection. (B) and (C) Quantification of cell cycle phases (G0, G1, S, and M) in CD317^-^ and CD317^+^ fractions of the Lin^-^ (B) and LK (C) cell populations from pIpC stimulated mice at 4 h and 24 h post-injection. (D) Representative FACS plots showing cell cycle analysis using DAPI and intracellular expression of Ki67 in CD317^-^ and CD317^+^ fractions of Lin^-^ (upper panel) and LK (lower panel) cell populations from the BM of LPS-stimulated mice at 2 h and 24 h post-injection. (E) and (F) Quantification of cell cycle phases (G0, G1, S, and M) in CD317^-^ and CD317^+^ fractions of the Lin^-^ (E) and LK (F) cell populations from pIpC-stimulated mice at 2 h and 24 h post-injection. Data are presented as average ± SD, a summary of three independent experiments, n≥3 mice per group; * - p<0.05, ** - p<0.01, *** - p<0.001.

